# Pharmacological manipulations of the dorsomedial and dorsolateral striatum during fear extinction have opposing effects on fear renewal

**DOI:** 10.1101/2024.01.17.576042

**Authors:** Margaret K Tanner, Alyssa A Hohorst, Jessica Westerman, Carolina Sanchez Mendoza, Rebecca Han, Nicolette A Moya, Jennifer Jaime, Lareina M Alvarez, Miles Dryden, Aleezah Balolia, Remla Abdul, Esteban C Loetz, Benjamin N Greenwood

**Affiliations:** Department of Psychology, University of Colorado Denver, Denver, CO; Department of Integrative Biology, University of Colorado Denver, Denver, CO; Department of Neuroscience, Northwestern Feinberg School of Medicine, Chicago, IL; The Neuroscience Graduate Program, University of Michigan, Ann Arbor, MI 48109, USA; Neuroscience Program, University of Colorado Anschutz Medical Campus, Denver, CO

## Abstract

Systemic manipulations that enhance dopamine (DA) transmission around the time of fear extinction can strengthen fear extinction and reduce conditioned fear relapse. Prior studies investigating the brain regions where DA augments fear extinction focus on targets of mesolimbic and mesocortical DA systems originating in the ventral tegmental area, given the role of these DA neurons in prediction error. The dorsal striatum (DS), a primary target of the nigrostriatal DA system originating in the substantia nigra (SN), is implicated in behaviors beyond its canonical role in movement, such as reward and punishment, goal-directed action, and stimulus-response associations, but whether DS DA contributes to fear extinction is unknown. We have observed that chemogenetic stimulation of SN DA neurons during fear extinction prevents the return of fear in contexts different from the extinction context, a form of relapse called renewal. This effect of SN DA stimulation is mimicked by a DA D1 receptor (D1R) agonist injected into the DS, thus implicating DS DA in fear extinction. Different DS subregions subserve unique functions of the DS, but it is unclear where in the DS D1R agonist acts during fear extinction to reduce renewal. Furthermore, although fear extinction increases neural activity in DS subregions, whether neural activity in DS subregions is causally involved in fear extinction is unknown. To explore the role of DS subregions in fear extinction, adult, male Long-Evans rats received microinjections of either the D1R agonist SKF38393 or a cocktail consisting of GABA_A_/GABA_B_ receptor agonists muscimol/baclofen selectively into either dorsomedial (DMS) or dorsolateral (DLS) DS subregions immediately prior to fear extinction, and extinction retention and renewal were subsequently assessed drug-free. While increasing D1R signaling in the DMS during fear extinction did not impact fear extinction retention or renewal, DMS inactivation reduced later renewal. In contrast, DLS inactivation had no effect on fear extinction retention or renewal but increasing D1R signaling in the DLS during extinction reduced fear renewal. These data suggest that DMS and DLS activity during fear extinction can have opposing effects on later fear renewal, with the DMS promoting renewal and the DLS opposing renewal. Mechanisms through which the DS could influence the contextual gating of fear extinction are discussed.

**Highlights:**

- Dorsolateral striatum D1 receptor signaling during fear extinction reduces renewal
- Neural activity in the dorsomedial striatum during fear extinction permits renewal
- Dorsal striatum subregions have opposing roles in contextual gating of fear extinction

**Graphical Abstract:** 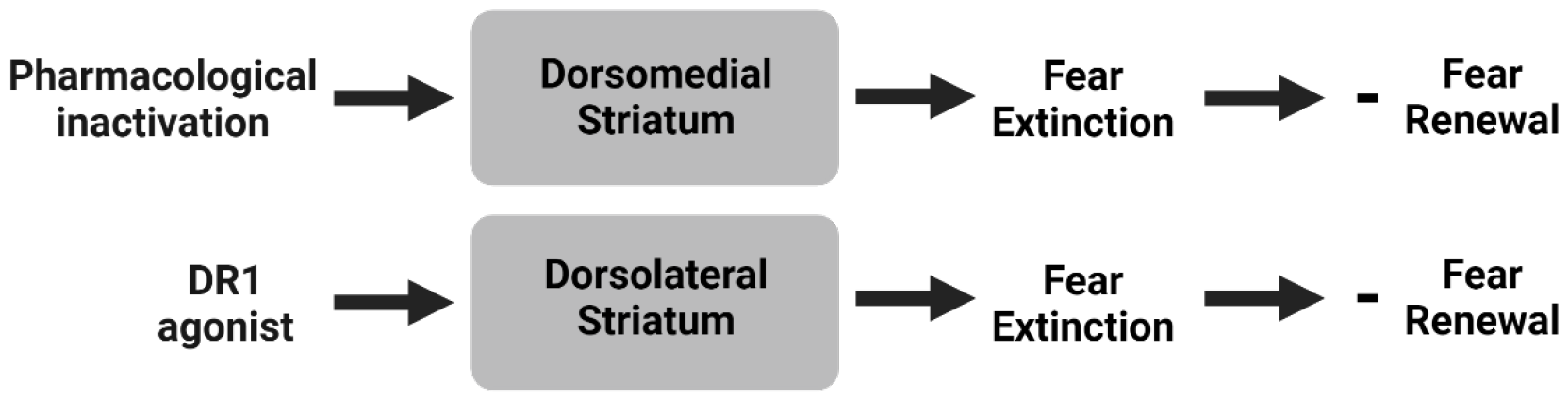

## 1. Introduction

Fear conditioning is an associative learning process during which an initially neutral stimulus (conditioned stimulus; CS) comes to evoke a fear response after its association with an aversive event (unconditioned stimulus; US). Excessive fear learning and impaired inhibition of fear memories contribute to anxiety- and trauma-related disorders, such as post-traumatic stress disorder. Extinction-based exposure therapy is a commonly used behavioral therapy to treat the intrusive fear responses present in anxiety and trauma-related disorders. During fear extinction, repeated presentations of the fear-evoking CS in the absence of the aversive US causes a reduction in the fear response associated with the CS [1]. However, fear extinction memories are labile and susceptible to memory interference phenomena which contribute to relapse of conditioned fear after extinction [2]. One such relapse phenomena is fear renewal, during which conditioned fear returns after extinction in contexts different from the context in which extinction took place [2-4]. Since relapse of previously extinguished fear limits the long-term success of exposure therapy, research efforts focus on identifying strategies to enhance fear extinction and reduce relapse.

Dopamine (DA) is emerging as a key player in fear extinction [5, 6] and is a potential target to reduce relapse of fear after extinction. When administered after fear extinction, during the memory consolidation phase, systemic DA agonists are consistently reported to enhance fear extinction retention [6-9]. Importantly, systemic DA agonists such D1/D5 receptor agonists [7] and L-DOPA [8, 9] attenuates the contextual modulation of fear extinction memory, thereby reducing fear renewal. Together, these data indicate that fear extinction augmented by DA manipulations can be resistant to relapse.

Studies investigating the role of specific brain regions where DA acts to modulate fear extinction have focused on targets of midbrain DA neurons originating in the ventral tegmental area (VTA), such as the nucleus accumbens (NAc), basolateral amygdala (BLA), prefrontal cortex (PFC), and hippocampus, given the preponderance of evidence for critical roles of these regions in fear extinction [10, 11]. VTA DA neurons projecting to the NAc are thought to contribute to fear extinction by encoding the prediction error that occurs when the expected US no longer follows the CS during fear extinction [12-14]. In the BLA, DA receptor antagonists prior to [15], but not after [16], fear extinction interfere with fear extinction retention, suggesting that DA in the BLA contributes to the acquisition of fear extinction. Some data suggest that DA receptor signaling in the infralimbic cortex of the PFC contributes to the consolidation of fear extinction memory [6, 16, 17], while other data indicate VTA DA projections to the PFC oppose fear extinction and promote relapse [14, 18]. In the hippocampus, DA D1 receptor (D1R) signaling has been reported to enhance the consolidation of contextual fear extinction [19]. Although DA in targets of the VTA can support the acquisition and/or consolidation of fear extinction, whether fear extinction augmented by DA in these regions is resistant to relapse has not been thoroughly explored. This is a critical omission, as manipulations that enhance fear extinction do not necessarily reduce relapse [20-23]. The brain region(s) where DA agonists act to reduce relapse following fear extinction thus remain unknown. Also unknown is whether DA in targets of midbrain DA neurons other than the NAc, BLA, PFC, and hippocampus also contribute to fear extinction.

The dorsal striatum (DS) is a primary target of midbrain DA neurons arising from the substantia nigra (SN). The DS is increasingly implicated in behaviors beyond its canonical role in movement, including reward and punishment [24-26], emotional behavior [27], goal-directed action [28], stimulus-response learning [29], and aversive learning and memory such as fear conditioning [30, 31]. The DS, therefore, is another region where DA could potentially influence fear extinction, yet few studies have investigated the role of the DS in fear extinction.

We have previously reported that stimulation of SN DA neurons in rats during fear extinction with designer receptors exclusively activated by designer drugs (DREADDs) had no impact on freezing during extinction acquisition, but enhanced fear extinction retention and completely prevented fear renewal in a novel context [32]. Although this study did not clarify the SN target sites mediating the effects of SN DA stimulation, expression of terminal DREADDs was found to be greatest in the DS, and the effect of SN DA stimulation during extinction on later fear renewal was mimicked by injection of a D1R agonist into the DS immediately prior to extinction [32]. These data point to a role for DA neurons in the SN and DS D1R signaling in augmenting fear extinction such that fear extinction becomes resistant to relapse. However, D1R agonists in this study were injected into the middle of the DS, so it remains unknown where in the DS the drugs were exerting their behavioral effects. This is an important overlooked question, since the DS is subdivided into subdivisions which subserve unique functions of the DS: the dorsomedial (DMS) and dorsolateral (DLS) subregions [33, 34]. Additionally, potential roles of the DMS and DLS in supporting “normal” fear extinction processes (e.g., fear extinction in the absence of D1R agonists) are not established.

The current studies investigated the roles of DS subregions and D1R signaling within DS subregions in cued fear extinction and renewal. To determine the DS subregion mediating the previously reported ability of intra-DS D1R agonist to reduce later fear renewal [32], the D1R agonist SKF38393 was injected selectively into the DMS or DLS, immediately prior to fear extinction. Next, the DMS and DLS were temporarily inactivated during fear extinction with injections of the GABA_A_/GABA_B_ receptor agonists muscimol/baclofen to determine whether these DS subregions contribute causally to normal fear extinction. Results suggest that DS subregions have opposing influences on the contextual gating of fear extinction memory. The DMS, but not DLS, contributes to normal fear extinction by permitting fear extinction memory to remain susceptible to fear renewal. In contrast, increasing D1R signaling in the DLS, but not DMS, during fear extinction reduces later renewal while having no impact on fear extinction acquisition or retention. Potential mechanisms through which the DS could influence contextual gating of fear extinction are discussed.

## 2. Materials and Methods

### 2.1. Animals and housing

A total of 100 adult (postnatal day 56 on arrival), male, Long-Evans rats were purchased from Charles River. Rats were pair-housed in Plexiglas cages (24 L x 45.4 W x 21 H cm) and kept on a 12 h light-dark cycle (6:00 am – 6:00 pm LD) with *ad libitum* access to food and water. For the duration of the experiment, rats were housed in a temperature- (22° C) and humidity-controlled vivarium accredited by the Association for Assessment and Accreditation of Laboratory Animal Care located on the University of Colorado Denver Auraria campus.

Experimental procedures took place during the light cycle between 9:00 am – 12:00 pm. All experimental protocols were approved by the University of Colorado Denver institutional animal care and use committee.

### 2.2. Stereotaxic surgery

Bilateral 26-gauge guide cannulae (Plastics One, Roanoke, VA, USA) were targeted to either the DMS (+0.5 mm anterior, ± 1.8 mm lateral, − 4.6 mm ventral from the top of the skull) or DLS (+0.5 mm anterior, ± 4.2 mm lateral, − 4.9 mm ventral from the top of the skull) according to the Paxinos and Watson rat brain atlas [35] and our recent work [36]. All surgeries were performed under ketamine (75.0 mg/kg i.p.) and medetomidine (0.5 mg/kg i.p.) anesthesia. Atipamezole (0.5 mg/kg i.p.) was given as a reversing agent to speed recovery. Animals received injections of carprofen (5 mg/kg) and penicillin G (22,000 IU/rat) subcutaneously at induction and every 24 h for 72 h post-surgery. Before experimental procedures began rats were allowed to recover for 1 week. Rats were handled daily after surgery during which time dummy cannulae were removed from the guide cannulae and replaced to maintain the patency of the lumen. Brain tissue was inspected under a microscope (Olympus BX53; Center Valley, PA) to verify cannulae placements after the conclusion of behavioral procedures. Rats with misplaced cannulae were excluded from the experiment.

### 2.3. Drugs

D1R agonist 2,3,4,5,-tetrahydro-7,8-dihydroxy-1-phenyl-1*H*-3-benzazepine hydrochloride (<u>SKF38393</u>; Sigma-Aldrich, St Louis, MO) was dissolved in sterile saline at a concentration of 0.5 μg/μL, based on prior work [32, 37]. Muscimol (Musc; 0.03 nmol/uL; Thermo Fisher Scientific, Waltham, MA) and baclofen (Bac; 0.3 nmol/uL; Thermo Fisher Scientific) were dissolved together in sterile saline. Concentration of Musc/Bac was based on prior work administering Musc/Bac into DS subregions [36, 38].

### 2.4. Microinjections

Rats were moved to a room adjacent to the housing room, where dummy cannulae were replaced by microinjectors connected to infusion pumps (World Precision Instruments, Sarasota, FL) with PE50 tubing. Microinjectors extended 0.5mm below the guide cannulae tips. Freely moving rats received 1 μL/hemisphere of either saline, SKF38393, or Musc/Bac cocktail at a rate of 0.5 μL/min. Microinjectors were left in place for 2 min after infusion to allow for diffusion. Microinjections occurred 15 min prior to fear extinction training, as this was the timepoint used in our prior studies using these drugs [32, 36]. All rats were exposed to mock infusion procedures in the infusion room once a day for 2 days prior to microinjections to habituate rats to the microinjection procedure. Prior to exclusion of rats with misplaced cannulae, n=15 / group were used for the D1R agonist study and n = 10 / group were used for the Musc/Bac study.

### 2.5. Behavioral testing

Auditory fear conditioning, extinction training, extinction memory retention, and renewal procedures were similar to our prior work [39]. Each test was separated by 24 h and occurred in distinct contexts (A, B, C design). A schematic depicting the design of the experiments is provided in Figure 1. Freezing, an innate fear response, was defined as the absence of all movement except that required for respiration and was used as the measure of fear in all behavior tests. Behavior was recorded with overhead cameras and videos were later scored for freezing during each CS both with Noldus EthoVision XT (Leesburg, VA) and by a human experimenter blind to treatment conditions of the animals. EthoVision was also used to quantify locomotor activity prior to the first CS during each behavioral test.

**Figure 1.**
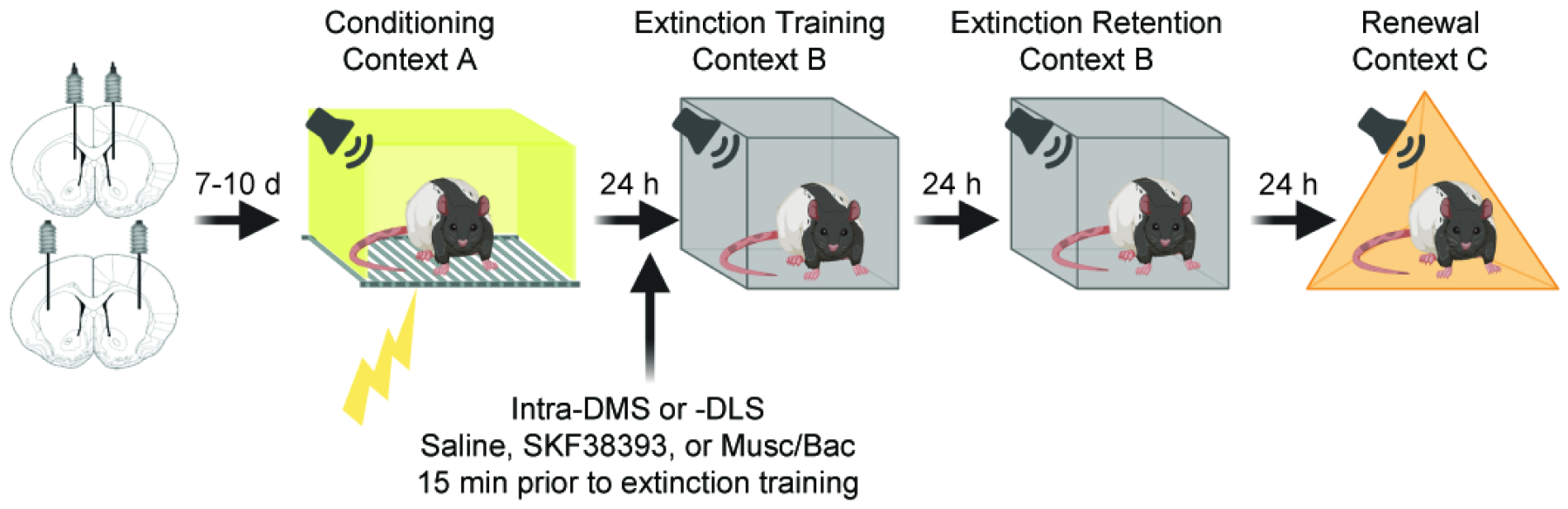
Experimental timeline. One week after stereotaxic implantation of bilateral cannulae into the dorsomedial (DMS) or dorsolateral (DLS) striatum, rats were exposed to auditory fear conditioning in Context A (yellow). The next day, rats received bilateral infusions of either sterile saline, the D1R agonist SKF38393, or a cocktail of the GABA_A_/GABA_B_ receptor agonists Muscimol/Baclofen (Musc/Bac) through the cannulae 15 min prior to fear extinction training (arrow) in Context B (gray). Fear extinction memory retention was tested 24 h later in the extinction context, Context B. Twenty-four hours later, rats were re-exposed to the extinguished CS in a novel Context C (orange) to assess fear renewal, which is the return of fear after extinction in a context different from where extinction occurred.

One week after surgery, rats were transported in their home cages to a behavioral testing room to undergo fear conditioning. Rats were placed into custom, rectangular conditioning chambers (Context A; 20″ W × 10″ D × 12″ H) with a shock grid floor (Coulbourn Instruments, Allentown, PA). Each conditioning chamber was housed inside individual sound-attenuating cabinets illuminated with red lights. Rats were allowed 3 min to explore the context, after which 4 auditory CS (10 sec, 80 dB, 2 kHz), each co-terminating with a 1-sec, 0.8-mA foot shock US, were delivered on a 1 min inter-trial interval. Auditory stimuli and foot shocks were delivered through Coulbourn tone generators and shock scramblers controlled via Noldus EthoVision XT software through a custom interface. All rats remained in the conditioning chamber, Context A, for 1 min after the last shock before being transported back to their housing room. Chambers were cleaned with water between rats.

Auditory fear extinction training took place in Context B, 24 h after fear conditioning. Rats were assigned to Saline and Drug groups based on freezing levels during fear conditioning to avoid arbitrary differences in conditioned fear acquisition between groups. Fifteen min after microinjection, rats were placed into a custom Plexiglass chamber that was either rectangular (15″ W × 15″ D × 20″ H) with a textured floor or a triangle (15″ sides × 20″ H) with a smooth floor. Rectangular and triangular chambers were counterbalanced so half the rats were exposed to fear extinction training in the triangular chambers and the other half in the rectangular chambers. Context B chambers were housed in the same sound-attenuating cabinets used for conditioning, but all other contextual features and discrete cues differed between contexts A and B. Rats were transported to the sound-attenuating cabinets in the testing room in their assigned Context B chambers. The sound-attenuating cabinets included vanilla scent and were illuminated with bright white lights, while a fan located in the cabinet was used to provide ventilation and background noise. After a 3 min exploration period, the auditory CS was presented 20 times (1 min ITI) in the absence of the foot shock US. Rats were removed from Context B 1 min after the last CS presentation and returned to their home cages. Context B was cleaned with 10% ethanol between tests. The auditory fear extinction procedure was repeated the next day, without microinjections, as a measure of fear extinction memory retention.

Fear renewal testing occurred 24 h following the fear extinction memory retention test in a novel Context C. Rats that underwent extinction in the rectangular Plexiglas chamber were now placed into the triangular chamber, and vice versa. Rats were transported in their Context C chambers to the sound-attenuating cabinets, which contained a raspberry scent and were dimly lit by a lamp located outside of the cabinets. Context C chambers were cleaned with 1% acetic acid between tests. After a 3 min exploration period, the auditory CS was presented 5 times (1 min ITI) in the absence of the foot shock US. We have previously reported that very little fear transfers from Context B to Context C [32, 40], indicating that rats perceive these contexts as unique.

### 2.6. Statistical Analyses

Freezing scores during CS presentations obtained from an experimenter blind to treatment condition of the animals were averaged with scores from Noldus EthoVision XT. Freezing prior to the first CS presentation in all contexts was negligible and is not shown. Group differences in freezing across all 4 CS-US pairings during fear conditioning were analyzed with repeated measures ANOVA. Freezing during fear extinction and fear extinction memory retention tests were collapsed into 4 blocks consisting of 5 CS each and group differences were analyzed with repeated measures ANOVA. Shapiro–Wilk and Brown-Forsythe tests verified normality and equal variance of the data, respectively, prior to running ANOVAs. Freezing during all 5 CS exposures during fear renewal were averaged and group differences were compared with an unpaired t-test. Unpaired t-tests were used to compare group differences in locomotor activity prior to the first CS exposure during each behavioral test. Group differences were considered significant when p < 0.05.

## 3. Results

### 3.1. Increasing D1R signaling in the DLS, but not DMS, during fear extinction training reduces fear renewal

Prior work indicates that D1R agonists infused into the middle of the DS just prior to fear extinction reduce later fear renewal [32], but the DS subregion in which D1R signaling is acting to reduce fear renewal remains unknown. Here we administered SKF38393 selectively into either the DMS or DLS prior to fear extinction training to determine the DS subregion in which D1R agonist during fear extinction impacts later fear renewal. Location of cannulae tips in the DMS and DLS are shown in Figures 2A and 2G, respectively. After exclusion of rats with misplaced cannulae, final group sizes were: DMS Saline = 12; DMS SKF38393 = 14; DLS Saline = 15; DLS SKF38393 = 15.

**Figure 2.**
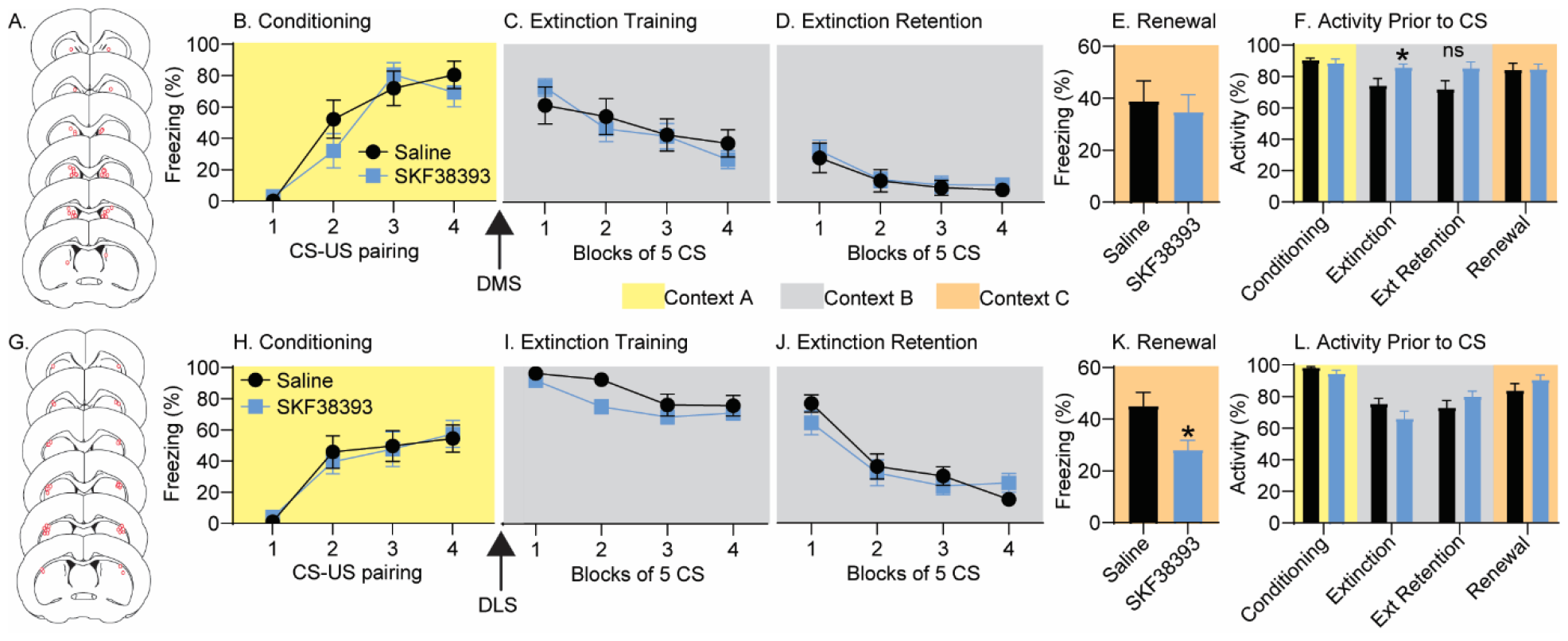
Behavioral effects of dopamine D1 receptor agonist SKF38393. Rats with bilateral cannulae aimed at the DMS (red dots; A) were exposed to auditory fear conditioning in Context A (yellow box) and freezing was measured (B). Rats received intra-DMS Saline or SKF38393 infusions 15 min prior to auditory fear extinction (arrow) in Context B (gray box; C). Fear extinction retention was measured the next day in Context B (D), followed 24h later by fear renewal in a novel Context C (orange box; E). Locomotor activity was quantified prior to the first CS exposure during each behavioral test (F). Intra-DMS SKF38393 increased locomotor activity in Context B prior to fear extinction (*p = 0.02). The group difference in activity prior to fear extinction retention was not significant (ns). Rats with bilateral cannulae aimed at the DLS (red dots; G) were exposed to auditory fear conditioning in Context A and freezing was measured (H). Rats received intra-DLS Saline or SKF38393 infusions 15 min prior to auditory fear extinction in Context B (I). Fear extinction retention was measured 24h later in Context B (J). The next day, rats were exposed to a novel Context C for fear renewal testing, during which prior intra-DLS SKF38393 reduced freezing relative to Saline (*p = 0.01; K). Locomotor activity was quantified prior to the first CS during each behavioral test (L). All freezing data represent group means ± SEM.

Rats with cannulae in the DMS all acquired auditory fear conditioning (*F*_(3, 72)_ = 43.2; p < 0.0001; Figure 2B) and there were no differences between rats subsequently assigned to Saline or SKF38393 groups (*F*_(1, 24)_ = 0.36; p = 0.6). Freezing decreased over time during extinction (*F*_(3, 72)_ = 15.16; p < 0.0001; Figure 2C) and there was no effect of SKF38393 on freezing during extinction training (*F*_(1, 24)_ = 0.03; p = 0.9), extinction memory retention (*F*_(1, 24)_ = 0.12; p = 0.7; Figure 2D), or fear renewal (*F*_(1, 24)_ = 0.16; p = 0.69; Figure 2E). SKF38393 injections into the DMS increased locomotor activity relative to Saline during the 3 min exploration period prior to the first CS during fear extinction (unpaired t test: p = 0.02), but not during any other behavioral test (Figure 2F).

Rats with cannulae in the DLS all acquired auditory fear conditioning (*F*_(3, 84)_ = 25.72; p < 0.0001; Figure 2H) and there were no differences between rats subsequently assigned to Saline or SKF38393 groups (*F*_(1, 28)_ = 0.005; p = 0.95). Freezing decreased over time during extinction (*F*_(3, 84)_ = 15.63; p < 0.0001; Figure 2I) and there was no effect of SKF38393 on freezing during extinction training (*F*_(1, 28)_ = 3.05; p = 0.09) or extinction memory retention (*F*_(1, 28)_ = 0.21; p = 0.65; Figure 2J). However, prior intra-DLS SKF38393 did reduce freezing in the novel Context C during the fear renewal test (unpaired t test: p = 0.01; Figure 2K). Intra-DLS SKF38393 had no effect on locomotor activity prior to the first CS during any behavioral test (Figure 2L).

### 3.2. Inactivation of the DMS, but not DLS, during fear extinction training reduces fear renewal

Involvement of DS subregions in fear extinction and relapse was investigated by temporary pharmacological inactivation. Saline or a cocktail of Musc/Bac was injected into either the DMS or DLS 15 min prior to fear extinction training, and fear extinction retention and renewal were subsequently assessed (Figure 1). Location of cannulae tips in the DMS and DLS are shown in Figures 3A and 3G, respectively. After exclusion of rats with misplaced cannulae, final group sizes were as follows: DMS Saline = 9; DMS Musc/Bac = 10; DLS Saline = 8; DLS Musc/Bac = 7.

**Figure 3.**
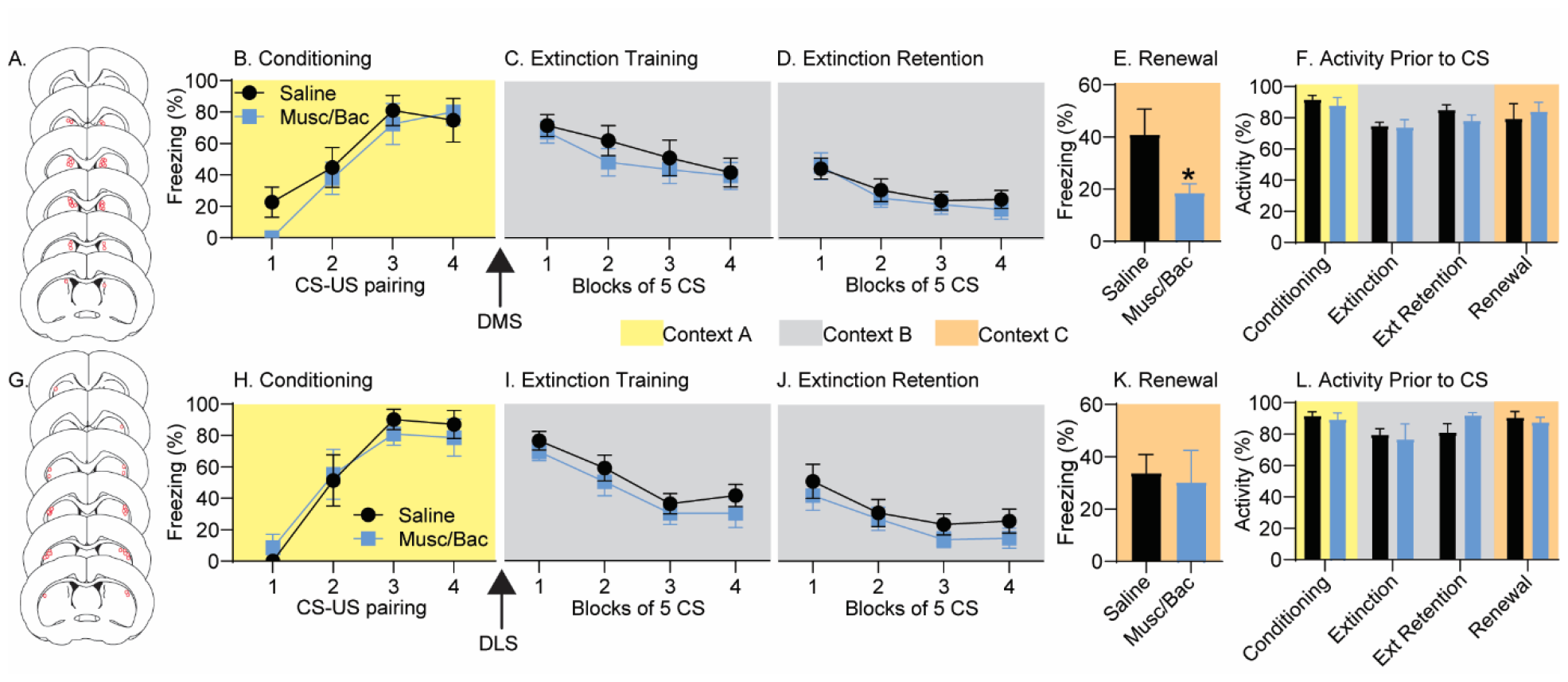
Behavioral effects of dorsomedial striatum (DMS) or dorsolateral striatum (DLS) inactivation. Rats with bilateral cannulae aimed at the DMS (red dots; A) were exposed to auditory fear conditioning in Context A (yellow box) and freezing was measured (B). Rats received intra-DMS Saline or a cocktail of GABA_A_/GABA_B_ receptor agonists Muscimol/Baclofen (Musc/Bac) 15min prior to auditory fear extinction (arrow) in Context B (gray box; C). Fear extinction retention was measured 24h later in Context B (D). The next day, rats were exposed to a novel Context C (orange box) for fear renewal testing, during which prior intra-DMS Musc/Bac reduced freezing relative to Saline (*p = 0.03; E). Locomotor activity was quantified prior to the first CS during each behavioral test (F). Rats with bilateral cannulae aimed at the DLS (red dots; G) were exposed to auditory fear conditioning in Context A and freezing was measured (H). Rats received intra-DLS Saline or Musc/Bac 15min prior to auditory fear extinction (arrow) in Context B (I). Fear extinction retention was measured 24h later in Context B (J), followed 24h later by fear renewal in a novel Context C (K). Locomotor activity was quantified prior to the first CS during each behavioral test (L). All freezing data represent group means ± SEM.

Rats with cannulae in the DMS all acquired auditory fear conditioning (*F*_(3, 51)_ = 22.36; p < 0.0001; Figure 3B) and there were no differences between rats subsequently assigned to Saline or Musc/Bac groups (*F*_(1, 17)_ = 0.83; p = 0.4). Freezing decreased over time during extinction (*F*_(3, 31)_ = 13.9; p < 0.0001) and there was no effect of Musc/Bac on freezing during either extinction training (*F*_(1, 17)_ = 0.38; p = 0.5; Figure 3C) or the fear extinction retention test 24 h later (*F*_(1, 17)_ = 0.13; p = 0.7; Figure 3D). However, rats that received Musc/Bac prior to fear extinction displayed reduced freezing in the novel context C during the fear renewal test (*F*_(1, 17)_ = 5.1; p = 0.03; Figure 3E). No group differences in locomotor activity prior to the first CS in any of the behavioral tests were observed (Figure 3F).

Rats with cannulae in the DLS all acquired auditory fear conditioning (*F*_(3, 39)_ = 23.8; p < 0.0001; Figure 3H) and there were no differences between rats subsequently assigned to Saline or Musc/Bac groups (*F*_(1, 13)_ = 0.01; p = 0.9). Freezing decreased over time during extinction (*F*_(3, 39)_ = 24.1; p < 0.0001) and Musc/Bac did not significantly alter freezing during extinction training (*F*_(1, 13)_ = 1.0; p = 0.3; Figure 3I), extinction retention (*F*_(1, 13)_ = 0.64; p = 0.4; Figure 3J), or fear renewal testing (*F*_(1, 13)_ = 0.06; p = 0.8; Figure 3K). No group differences in locomotor activity prior to the first CS during any behavioral test were observed (Figure 3L).

## 4. Discussion

The DS is increasingly implicated in behaviors beyond its canonical role in movement, but whether the DS contributes to fear extinction is unknown. We have previously reported that fear extinction increases neural activity in both the DMS and DLS, and that increasing D1R signaling in the DS during fear extinction reduces later fear renewal [32]. Here, we expand on these prior observations by investigating 1) the DS subregion in which increasing D1R signaling during fear extinction reduces later renewal, and 2) the causal role of DS subregion neural activity during fear extinction in later fear extinction retention and renewal.

When injected into the DMS just prior to fear extinction training, the D1R agonist SKF38393 had no impact on freezing during fear extinction acquisition, fear extinction memory retention, or fear renewal (Figure 2D-E). The lack of effect on freezing is unlikely to be caused by an insufficient drug dose, because intra-DMS SKF38393, relative to Saline, did increase locomotor activity in the extinction context prior to the first CS during fear extinction training (Figure 2F). These data suggest that the DMS is an unlikely target mediating the previously reported ability of intra-DS D1R agonist to reduce fear renewal [32]. We have previously observed that fear extinction increases *cfos* mRNA selectively in D1R-expressing neurons in the DMS [32]. The fact that DMS D1R neurons are already recruited during normal fear extinction could explain why a D1R agonist has no additional effect in this region.

In contrast to its lack of effect when injected into the DMS, SKF38393 injected into the DLS prior to fear extinction training reduced freezing during the fear renewal test, relative to intra-DLS saline (Figure 2K). The observation that intra-DLS SKF38393 had no impact on locomotor activity (Figure 2L) reveals that the reduction in freezing during renewal is not due to a general increase in locomotor activity elicited by SKF38393. The reduction in fear renewal caused by intra-DLS SKF38393 also seems not to be a consequence of enhanced fear extinction acquisition or retention, as no effects of SKF38393 on freezing during extinction training or retention were observed (Figures 2I and 2J). This pattern of data is identical to the previously observed effect of intra-DS SKF38393, which also reduced fear renewal without enhancing fear extinction retention in the fear extinction context [32]. These data suggest that increasing D1R signaling in the DLS during fear extinction somehow frees fear extinction memory from contextual modulation, thus reducing freezing in contexts different from the extinction context. The DLS could thus be a potential target mediating the ability of systemic DA agonists to similarly reduce contextual dependency of fear extinction memory [7-9].

It is of interest to consider how the current results inform the mechanisms through which SN DA stimulation during fear extinction reduces later renewal. In contrast to the pattern of neural activity noted in the DMS during fear extinction, fear extinction selectively increases *cfos* in D2 receptor-expressing neurons in the DLS [32]. However, activity of DLS D1R-expressing neurons is increased when SN DA neurons are chemogenetically stimulated during fear extinction [32]. These prior data, combined with the current observation that D1R agonist administered during extinction reduces later renewal when injected into the DLS, but not DMS, support the possibility that SN DA stimulation during fear extinction reduces renewal by increasing activity of D1R neurons in the DLS. Unlike intra-DLS SKF38393, however, SN DA stimulation during fear extinction training also enhanced fear extinction retention [32]. It is possible that the enhancement of fear extinction retention and the reduction of fear renewal elicited by SN DA stimulation are not causally related but are instead mediated by unique SN targets. While DLS D1R neurons seem to be a likely target mediating the reduction in renewal caused by SN DA stimulation, the SN target supporting enhanced fear extinction has yet to be identified. It is unlikely to be the DMS, since intra-DMS D1R agonist failed to enhance fear extinction retention (Figure 2D).

Injecting D1R agonist into selective DS subregions reveals which DS subregion is a potential target whereby increasing D1R signaling could augment fear extinction and reduce relapse but does not divulge the potential roles of DS subregions in supporting “normal” fear extinction (e.g., fear extinction in the absence of D1R agonist). To determine whether neural activity within unique DS subregions is necessary for normal fear extinction, a cocktail of the GABA_A_/GABA_B_ receptor agonists Musc/Bac was injected directly into the DMS or DLS prior to fear extinction. Increasing confidence that Musc/Bac can successfully inhibit neural activity in DS subregions is the observation that the same Musc/Bac cocktail as used here inhibits neural activity in the DMS and DLS elicited by a bout of voluntary wheel running [36]. Similar to prior reports [36], neither DMS nor DLS inhibition altered general locomotor activity (Figures 3F and 3L). Thus, any observed effect of Musc/Bac on freezing is likely independent of alterations in general activity.

It is not particularly surprising that neither DMS nor DLS inactivation impacted fear extinction acquisition or retention (Figure 3), as neither DS sub-region has been reported to be critical to the Pavlovian associative learning process underlying fear extinction. Rather than contributing to fear extinction directly, neural activity observed in these regions during fear extinction could reflect context exploration and/or attempts to escape the context supported by goal-directed or stimulus-response strategies mediated by the DMS and DLS, respectively [29].

Interestingly, results indicate that DMS activity during fear extinction is necessary for fear extinction memory to remain susceptible to typical fear renewal (Figure 3E). Since DMS inactivation during fear extinction selectively altered the contextual modulation of fear extinction memory and not fear extinction *per se*, it seems likely that the DMS somehow contributes to contextual gating of fear extinction. Consistent with a primary role of the hippocampus in contextual processing, contextual modulation of fear extinction involves the hippocampus [41-43] and hippocampal projections to the amygdala [44, 45]. If the hippocampus is the driver of contextual modulation of fear extinction, then what is the role of the DMS? Contextual learning has been reported to involve cooperation between the DMS and the hippocampus in a variety of paradigms [46-51]. For example, disruption of DMS activity renders T-maze performance impervious to contextual cues and animals instead rely on hippocampal-independent, DLS-mediated “response” strategies for T-maze navigation [33, 48]. It is therefore possible that DMS inactivation during fear extinction could interfere with hippocampal-dependent contextual processing to render fear extinction memories free from contextual gating. Consistent with this hypothesis, the reduction in fear renewal produced by stimulation of SN DA neurons during extinction is associated with altered cFos expression in the CA1 of the hippocampus [32], which receives projections from the DS [52]. Additional research will be necessary to determine how DMS activity during fear extinction contributes to fear renewal and whether a DMS-hippocampal interaction mediating contextual processing plays a role. Involvement of interactions between DS and other regions involved in fear extinction, such as the PFC and amygdala, should also be considered in future studies.

A similar theoretical framework could explain how increasing D1R signaling in the DLS during fear extinction likewise reduces contextual gating of extinction. Whereas the DMS could cooperate with the hippocampus during fear extinction to enable contextual modulation of fear extinction memory (and promote renewal), increasing DLS D1R signaling during extinction could reduce renewal by impairing contextual processing. This notion is consistent with accumulating evidence suggesting that the DMS and DLS compete for neural control of behavior. DLS inhibition, for example, can facilitate DMS [53, 54] and hippocampal [55]-dependent behavioral strategies, and vice-versa [33, 48]. It is therefore plausible that increasing D1R signaling in the DLS during fear extinction could inadvertently inhibit the DMS, thus mimicking the effects of intra-DMS Musc/Bac and reducing fear renewal.

It is relevant to consider situations in which DLS activity during fear extinction could diminish contextual gating of fear extinction and reduce fear renewal. Since DLS inactivation during fear extinction failed to alter later fear renewal, a role for the DLS seems to only be revealed by manipulations that increase DLS activity (and perhaps specifically D1R-expressing neural activity) during fear extinction. Intra-DLS D1R agonist and stimulation of SN DA neurons are two obvious manipulations that increase DLS D1R signaling and reduce renewal, but there are others. Female rats exposed to fear extinction training during estrous phases characterized by high levels of ovarian hormones (proestrus and estrus) do not display fear renewal [40], and stimulus-evoked DA release in the DLS is greater in female rats during these phases [56]. Additionally, a bout of physical activity during the acquisition or consolidation of fear extinction reduces fear renewal in both rodents [39, 40, 57] and humans [58-60]. Physical activity sensitizes DA release in the DLS [61] and potentiates stress-evoked activity of DLS D1R-expressing neurons [62]. The potential role of DLS D1R signaling in the mechanisms by which high ovarian hormones and exercise at the time of fear extinction reduce later renewal deserves further exploration.

In summary, DS sub-regions seem to have an opposing influence on fear renewal. DMS neural activity during fear extinction is necessary for normal fear renewal, while increasing DLS D1R signaling during fear extinction reduces fear renewal. Results implicate the DS in contextual modulation of cued fear extinction memory. Future work is required to identify how DS subregions are involved in the contextual gating of fear extinction, potentially through interactions with the hippocampus. Regardless of the mechanism, the DS should be considered as a potential target for the reduction of fear relapse after extinction.

## Acknowledgements

This work was funded by R15MH114026 awarded to BNG.

